# Associations Between Plasma Lipoprotein(a) Levels and Circulating Monocyte Subsets Differ across Populations

**DOI:** 10.64898/2026.06.01.729444

**Authors:** Alexander C. Bashore, Benedek Halmos, Anouk G. Groenen, Anastasiya Matveyenko, Nelsa Matienzo, Lucie Y Zhu, Sergio Mosquera Restrepo, Yihao Li, Hanrui Zhang, Marit Westerterp, Muredach Reilly, Gissette Reyes-Soffer

## Abstract

**Background:** Lipoprotein(a) [Lp(a)] is a causal risk factor for cardiovascular disease (CVD), with plasma concentrations higher in Black individuals than in White individuals. Published findings support a model in which high Lp(a), in part through oxidized phospholipid (oxPL)-mediated signaling, promotes a pro-inflammatory monocyte phenotype that may contribute to arterial wall inflammation and the development of CVD. Here, we examined the relationship between Lp(a) concentrations and isoform size and the distribution of circulating monocyte populations in Black and White individuals using single-cell RNA sequencing (scRNA-seq) data.

**Methods:** Using standardized assays, we measured plasma Lp(a) levels, isoform size, inflammatory markers, and oxidized phospholipids in stored plasma samples from our previously published cohort of 128 participants. After excluding smokers and individuals with type II diabetes, a total of 34 participants (20 Black participants, 14 White participants) were included in analyses. Participants were further stratified by plasma Lp(a) levels into normal Lp(a) [median 12.6 nmol/L, with 15 individuals (8 Black participants) and high Lp(a) (median 159 nmol/L, with 19 individuals (12 Black partcipants)]. Multivariable linear regression was used to assess the association between plasma Lp(a) levels, Lp(a)-oxPL, and the proportion of monocyte subsets. Across all participants, scRNA-seq data identified six classical monocyte subsets, one non-classical monocyte subset, one MHCII^hi^ monocyte subset, and one interferon (IFN)-responsive monocyte subset.

**Results:** The distribution of monocyte subsets was similar in individuals with normal versus high Lp(a) levels. Despite the small study cohort, self resported race modified the assocation between plasma Lp(a) levels and the proportion of non-classical monocytes (*P* = 0.032), which was inversely associated in White participants (*P* = 0.028), but not in Black partcipatns (*P* = 0.470). OxPL bound to APO(a) showed a positive correlation with plasma Lp(a) levels (R^2^ = 0.84; *P* = 3.3×10^−14^), and non-classical monocytes (*P*_Whites_ = 0.027; *P*_Blacks_ = 0.275; *P*_Interaction_ = 0.014). Race also modified the association between plasma Lp(a) levels and the proportion of classical 2 monocytes (*P* = 0.027).

**Conclusions:** These results underscore the importance of self reported race when analysing studies on Lp(a), monocytes and, cardiovascular related monocyte immune functions.

## Introduction

Lipoprotein (a) [Lp(a)] is a genetically determined lipoprotein and a causal risk factor for cardiovascular disease (CVD).^1–3^ Structurally, Lp(a) consists of an APOB100-containing lipoprotein covalently linked to the glycoprotein apolipoprotein(a) [APO(a)].^4,5^ APO(a) is synthesized in the liver and exists in multiple isoforms whose size is determined by the number of Kringle IV type 2 (KIV-2) repeats.^6–8^ These repeats vary widely among individuals, producing APO(a) proteins ranging from approximately 300 to 800 kDa.^9–11^ Importantly, APO(a) isoform size is inversely related to circulating Lp(a) concentrations,^12– 14^ with the KIV-2 polymorphism accounting for approximately 30–70% of the inter-individual variability in plasma Lp(a) concentrations. Consistent with this strong genetic control, Black indivduals generally exhibit higher circulating Lp(a) levels compared to other populations.^15,16^

Despite the well-established role of Lp(a) in CVD, the mechanisms through which it promotes CVD remain imcompletely understood. Increasing evidence suggests that Lp(a) constributes to vascular inflammation through its effects on innate immune cells, particularly monocytes. One proposed mediator of these effects is the oxidized phospholipids (oxPL) carried by Lp(a).^17^ Supporting this, in experimental studies of subjects with elevated Lp(a), radiolabeled autologous peripheral blood mononuclear cells (PBMCs) demonstrate increased accumulation within the arterial wall of the carotid artery and aorta, suggesting Lp(a)-enhanced monocyte recruitment into the arterial wall.^17^ Consistent with this observation, monocytes from individuals with high Lp(a) exhibit upregulation of interferon (IFN)-α and IFN-γ response pathways. Importantly, these inflammatory signatures were attenuated following treatment with an antisense oligonucleotide that lowers Lp(a) concentrations by approximately 50%, which is accompanied by a reduced capacity for monocyte transendothelial migration.^18^ Collectively, these findings support a model in which elevated Lp(a), in part through oxPL-mediated signaling, promotes a pro-inflammatory monocyte phenotype that may contribute to arterial wall inflammation and the development of CVD.

The emergence of single-cell technologies has substantially advanced our understanding of circulating monocyte heterogeneity and the relationship of distinct monocyte subsets to CVD risk states.^19^ Notably, racial differences in circulating monocyte supopulations have also been reported, paralleling the well-established racial variation observed in plasma Lp(a) concentrations.^19^ These observations raise the possibility that Lp(a) may be a driver of racial patterns in monocyte population distributions or that relationships between Lp(a) and monocytes may differ across demographic populations.

To explore this possibility, we analyzed a sub-cohort from our previously published dataset examining circulating monocyte subsets.^19^ Specifically, we evaluated whether race modifies the relationship between plasma Lp(a) concentrations and monocyte subset distributions. Our analysis revealed that race moderates the association between Lp(a) and both non-classical monocytes and a subset of classical monocytes. In particular, serum Lp(a) concentrations were inversely associated with the proportion of non-classical monocytes in White participants, whereas this relationship was not observed in Black participants.

## Methods

We performed a cross-sectional observational study with secondary analysis of an existing cohort, incorporating single-cell transcriptomic profiling. A subcohort of available data and plasma samples from the recently Instiitutional Review Board approved and published cohort that enrolled 128 participants was used.^19^ Participants were recruited via direct advertising through the Columbia University RecruitMe registry (https://recruit.cumc.columbia.edu/) and ResearchMatch.org, Craigslist, IRB-approved flyers on campus and community bulletin boards, newspaper advertising, and recruitment from the New York-Presbyterian Hospital Preventive Cardiology/Lipid clinic. The subcohort was obtained after removing any individuals with type II diabetes and smokers and focusing exclusively on White and Black participants (N=34).

### Biochemical Measurements

Blood was collected during the visit after a 12-hour fast. A complete blood count with differential (assay 4646089), lipid panel (assay 4262402), cholesterol (assay 2921462), and Hemoglobin A1C (assay 2921552) were measured at the Center for Advanced Laboratory Medicine (CALM) at Columbia University Medical Center, exactly as described previously.^19^ Plasma LDL cholesterol levels were estimated using the Friedewald formula. Plasma APOB levels were measured by a human enzyme-linked immunosorbent assay (ELISA), purchased from Mabtech (3715-HP-2), Inc. Cincinnati, OH. Lp(a) plasma concentrations were measured using the isoform-independent sandwich ELISA developed by the Northwest Lipid Metabolism and Diabetes Research Laboratory.^20^ This method is validated to accurately determine Lp(a) concentrations independently of KIV^2^ repeats of APO(a). In addition, gel electrophoresis was used to measure APO(a) isoform size. In brief, 250 µL of plasma was diluted to 100 ng of protein in 40 µL of saline and combined with reducing buffer. The sample was then loaded onto an agarose gel, ran overnight at 123V and 4°C, transferred to a nitrocellulose membrane, immunoblotted, and imaged using the ChemiDoc MP Imaging System. This determined the isoform size present in the samples by comparison to in-house standard (combined material containing six APO(a) isoforms-38,32, 24, 19, 15, and 12 KIV-2 repeats). The expression of each isoform was established using the Image Lab software, which calculated relative proportions of the two isoforms based on the intensity profile of each lane. This method has an intra-sample variability that does not exceed 15%. A weighted isoform size (*wIS*) measurement was calculated.^21^ *wIS* represents the weighted average of the two apo(a) isoform sizes (present in an individual, weighted by their relative expression levels.

A chemiluminescent sandwich ELISA was used to measure oxPL per unit of APO(a) captured in plasma.^22,23^ Briefly, plasma was added to plates coated overnight with the monoclonal antibody LPA4.^24^ OxPLs were detected with biotinylated E06 in relative light units (RLU)/APO(a). Using the same ELISA, oxPL per unit of APOB100 was assessed,^22,23^ however, the Lp(a) was not immunoprecipitated prior to the assay. Therefore, we cannot distinguish whether oxPL was bound to APOB100 on LDL or Lp(a).

### Single-Cell RNA-Sequencing

Details on PBMC isolation and scRNA-seq of the study samples have been extensively described elsewhere.^19^ In brief, PBMCs were isolated from fresh blood and monocytes were FACS sorted by gating for CD14^+^CD16^+^HLA-DR^+/-^Lin^−^ cells. scRNA-seq data was generated at the Single Cell Core Facility at the Columbia University JP Sulzberger Genome Center using the Chromium Single Cell Gene Expression system (10x genomics), according to the manufacturere’s protocol, with 5000 cells and 100M reads targeted per sample. Fastq files were processed by Cell Ranger 5.0.1 from 10x genomics to generate count matrices of unique molecular identifiers (UMIs). Clustering was performed on filtered UMI count matrics based on 2500 highly variable RNAs, leading to the identification of 9 monocyte clusters. For the current study we generated a Uniform Manifold Approximation and Projection (UMAP) visualization with the data from the 34 participants included in the analysis.

### Statistical Analysis

#### Clinical and demographic data

Normality of clinical data as well as race, sex, and age were tested with a Shapiro-Wilk test. Data are presented as mean with standard deviation or median with interquartile range (IQR) where appropriate. Differences between continuous variable in the normal Lp(a) and high Lp(a) group were assessed by students t-test or Mann-Whitney U test where appropriate. Differences in categorical data were assessed by χ^2^ test.

### Associations with proportion of monocyte subsets

Because monocyte subsets are reported as a percentage of total monocyte subsets (*i*.*e*. the sum of each subset is 100% for all individuals), the compositional data is represented in a simplex rather than Eucledian space. In order to bring the compositional data into Eucledian space, we performed centered log-ratio (CLR) transformation using the formula 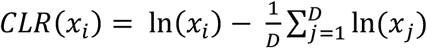, where x_i_ is the i-th component and D is the total number of components.

To investigate how race moderates the associations between Lp(a) and oxPL bound to APO(a) or APOB with the monocyte subset proportions, we performed multivariable linear regression with the CLR-transformed data as the outcome parameter. The resulting β-coefficient obtained from linear regression was transformed back to percentage change relative to the geometric mean using the formula % *C*h*ange* = (*e*^*B-coefficien*^ − 1) × 100. An interaction term for race and the predictor was used in multivariable linear regression analyses. We then report the race-specific effects for Black participants and White participants, as well as the *P*-value of the interaction term. A *P*-value <0.05 was considered statistically significant. We did not adjust for multiple comparisons as our compositional data is interdependent (*i*.*e*. any change in a given monocyte subset must be accompanied with changes in at least one other subset).

## Results

### Clinical characteristics

Plasma samples and associated clinical data were obtained from 34 participants within our previously published cohort,^19^ including 15 participants with normal (<50 nmol/L) and 19 participants with high (>50nmol/L) Lp(a) levels (**Supplemental Table I**). Median Lp(a) concentrations were 159.1nmol/L in the high Lp(a) group and 12.6 nmol/L in the normal Lp(a) group (**Table 1**). Compared with the normal Lp(a) group, participants with high Lp(a) were older (mean 45.7 *vs*. 37.6 years, *P* < 0.001), had higher total cholesterol (median 207 *vs*. 172 mg/dL, *P* = 0.023), low-density lipoprotein cholesterol (LDL-c) (median 131 *vs*. 84 mg/dL, *P* = 0.011), and oxidized phospholipids (oxPL) bound to APO(a) (median 33.1 *vs*. 3.59 RLU, *P* <0.001). As expected, the APO(a) wIS was smaller in participants with high Lp(a) compared to those with normal Lp(a) levels (21.3 *vs*. 26.8, *P <* 0.001). No significant differences were observed between groups in BMI, the number of circulating leukocytes, the proportions of immune cell types, or plasma levels of high-sensitivity C-reactive protein (hsCRP). Baseline characteristics of the study population are summarized in **Table 1**.

**Table 1.**
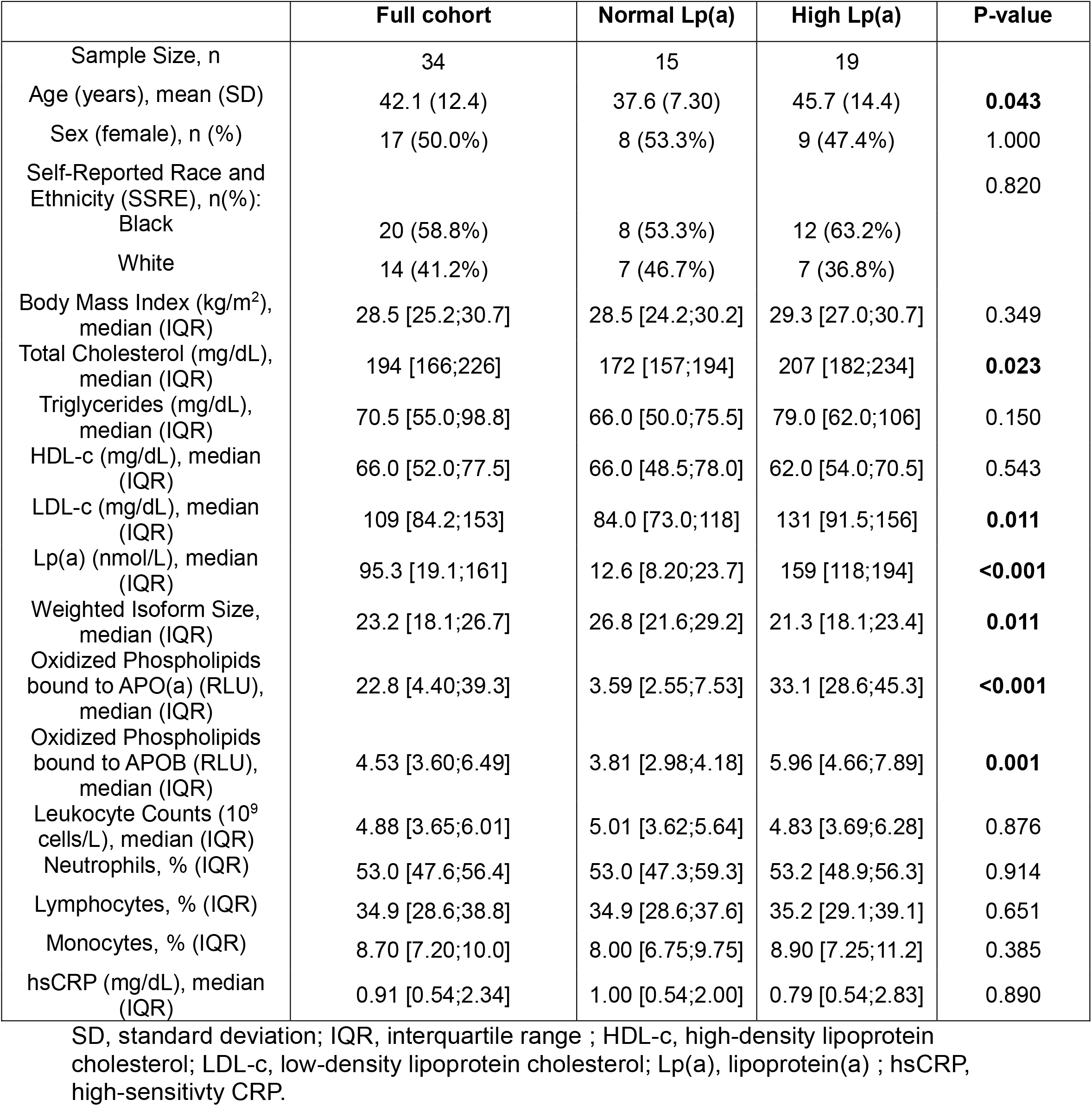
Characteristics of all 34 participants included in the analyses.

|  | Full cohort | Normal Lp(a) | High Lp(a) | P-value |
| --- | --- | --- | --- | --- |
| Sample Size, n | 34 | 15 | 19 |  |
| Age (years), mean (SD) | 42.1 (12.4) | 37.6 (7.30) | 45.7 (14.4) | **0.043** |
| Sex (female), n (%) | 17 (50.0%) | 8 (53.3%) | 9 (47.4%) | 1.000 |
| Self-Reported Race and Ethnicity (SSRE), n(%): |  |  |  | 0.820 |
| Black | 20 (58.8%) | 8 (53.3%) | 12 (63.2%) |  |
| White | 14 (41.2%) | 7 (46.7%) | 7 (36.8%) |  |
| Body Mass Index (kg/m<sup>2</sup>), median (IQR) | 28.5 [25.2;30.7] | 28.5 [24.2;30.2] | 29.3 [27.0;30.7] | 0.349 |
| Total Cholesterol (mg/dL), median (IQR) | 194 [166;226] | 172 [157;194] | 207 [182;234] | **0.023** |
| Triglycerides (mg/dL), median (IQR) | 70.5 [55.0;98.8] | 66.0 [50.0;75.5] | 79.0 [62.0;106] | 0.150 |
| HDL-c (mg/dL), median (IQR) | 66.0 [52.0;77.5] | 66.0 [48.5;78.0] | 62.0 [54.0;70.5] | 0.543 |
| LDL-c (mg/dL), median (IQR) | 109 [84.2;153] | 84.0 [73.0;118] | 131 [91.5;156] | **0.011** |
| Lp(a) (nmol/L), median (IQR) | 95.3 [19.1;161] | 12.6 [8.20;23.7] | 159 [118;194] | **<0.001** |
| Weighted Isoform Size, median (IQR) | 23.2 [18.1;26.7] | 26.8 [21.6;29.2] | 21.3 [18.1;23.4] | **0.011** |
| Oxidized Phospholipids bound to APO(a) (RLU), median (IQR) | 22.8 [4.40;39.3] | 3.59 [2.55;7.53] | 33.1 [28.6;45.3] | **<0.001** |
| Oxidized Phospholipids bound to APOB (RLU), median (IQR) | 4.53 [3.60;6.49] | 3.81 [2.98;4.18] | 5.96 [4.66;7.89] | **0.001** |
| Leukocyte Counts (10<sup>9</sup> cells/L), median (IQR) | 4.88 [3.65;6.01] | 5.01 [3.62;5.64] | 4.83 [3.69;6.28] | 0.876 |
| Neutrophils, % (IQR) | 53.0 [47.6;56.4] | 53.0 [47.3;59.3] | 53.2 [48.9;56.3] | 0.914 |
| Lymphocytes, % (IQR) | 34.9 [28.6;38.8] | 34.9 [28.6;37.6] | 35.2 [29.1;39.1] | 0.651 |
| Monocytes, % (IQR) | 8.70 [7.20;10.0] | 8.00 [6.75;9.75] | 8.90 [7.25;11.2] | 0.385 |
| hsCRP (mg/dL), median (IQR) | 0.91 [0.54;2.34] | 1.00 [0.54;2.00] | 0.79 [0.54;2.83] | 0.890 |

SD, standard deviation; IQR, interquartile range ; HDL-c, high-density lipoprotein cholesterol; LDL-c, low-density lipoprotein cholesterol; Lp(a), lipoprotein(a) ; hsCRP, high-sensitivity CRP.

### Distribution of monocyte subsets shows difference by race but not by Lp(a) levels

Using our previously published scRNA-seq data of circulating monocytes,^19^ we analyzed the data to investigate the relationship between Lp(a) levels and monocyte subset composition among the 34 participants in this substudy (**Supplemental Table III, Figure 1A**). Integration of the Uniform Manifold Approimiation and Projection (UMAP) visualization generated from the single-cell data enabled the identification of nine distinct monocyte populations (**Figure 1B**). Differential expression (DE) analysis confirmed the transcriptional signatures defining these including 6 classical monocyte subsets characterized by high CD99 expression, MHCII^hi^ monocytes with high HLA gene expression, interferon (IFN)-responsive monocytes enriched for interferon-stimulated genes such as ISG15, IFI6, and IFI44L, and non-classical monocytes defined by high FCGR3A (CD16) expression.^19^ When participants were stratified by normal and high Lp(a), UMAP visualization did not reveal any apparent differences in the overall distribution of monocyte populations between the two groups (**Figure 1D,E,F**).

**Figure 1.**
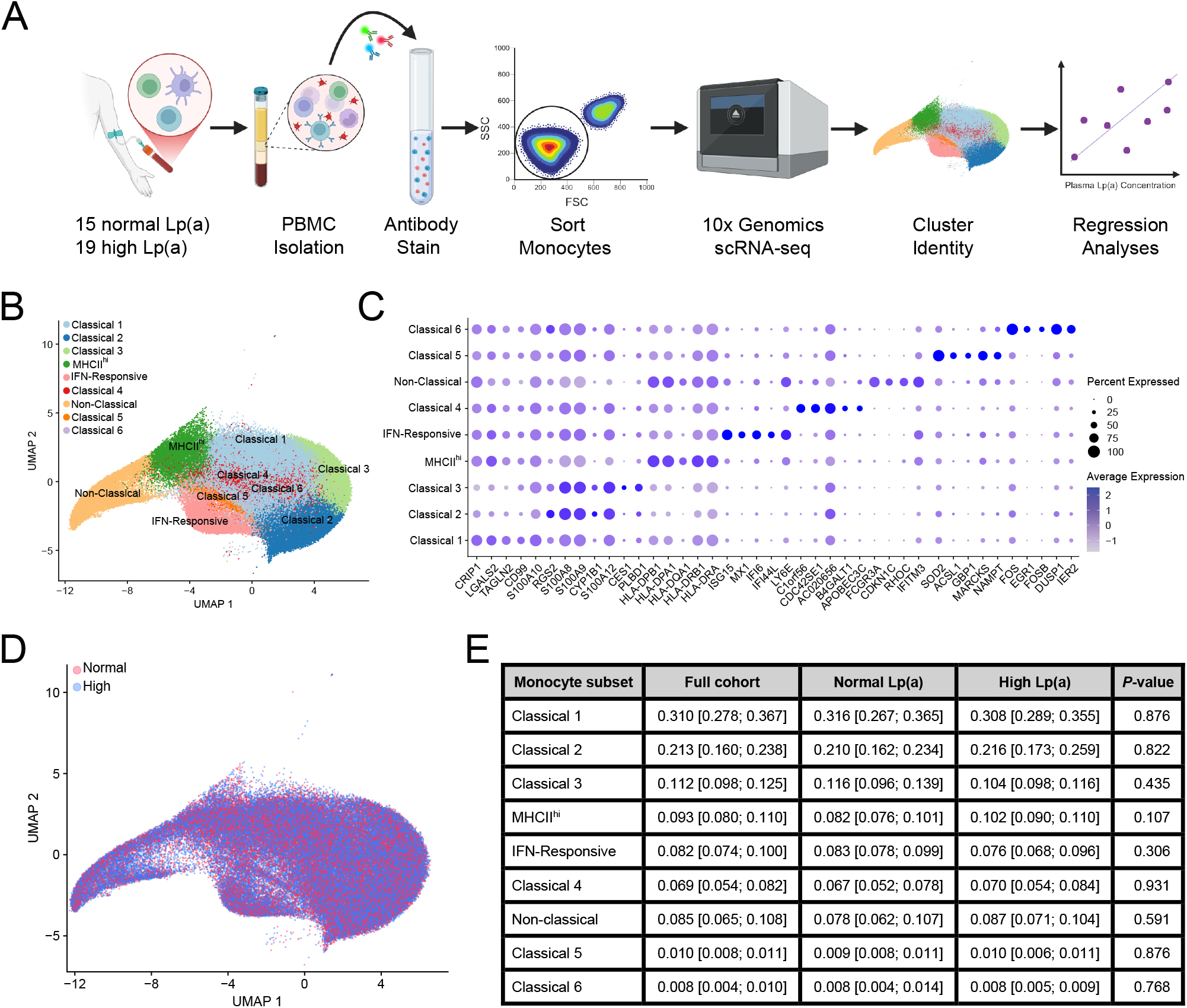
Single-cell RNA sequencing shows similar distribution of monocyte subsets in individuals with normal and high plasma Lp(a) levels. (**A**) Study design. (**B**) Uniform manifold approximation and projection (UMAP) visualization of intergrated data with cluster identity overlayed (n=34). (**C**) Dotplots displaying the top genes for each monocyte subset. (**D**) UMAP visualization of integrated data with plasma Lp(a) category overlayed. (**E**) Table showing median proportion with interquartile range (IQR) of every identified monocyte subset in the full cohort (n=34), as well as stratified for normal and high plasma Lp(a) categories (n=15 and n=19, respectively).

### Race modifies the association between Lp(a) concentrations and circulating monocyte subsets

Although no differences in monocyte subset proportions were observed between the high and normal Lp(a) groups when all participants were analyzed together, published data from the full cohort demonstrated racial differences in circulating monocyte subset distribution between Whites and Black individuals.^19^ Consistent with these findings, analysis of the present subcohort (n=34) also revealed racial differences in monocyte composition. White participants exhibited a higher proportion of MHCII^hi^ monocytes and lower proportion of non-classical monocytes compared to Black participants (Supplemental Figure I). In addition, White participants had higher proportions of Classical 1 and Classical 3 monocytes (Supplemental Figure I).

Since both plasma Lp(a) concentrations and monocyte subset distributions differ by race, we next assessed whether race modifies the relationship between Lp(a) and circulating monocyte subsets. Linear regression models incorporating an interaction term between race and high Lp(a) suggested that the association between high Lp(a) and classical 6 monocytes tended to be modified by race (*P*^interaction^ = 0.071; Supplemental Table II). Stratified analyses showed that Black participants with high Lp(a) had an 85% higher proportion of classical 6 monocytes compared with the partcipants with normal Lp(a) after adjustment for *wIS* and sex (*P* = 0.037; Figure 2; Supplemental Table II). In contrast, no difference in classical 6 monocyte proportions was observed in White participants with high or normal plasma Lp(a) levels. No race-specific effects were observed for the other monocyte subets (Supplemental Table II).

**Figure 2.**
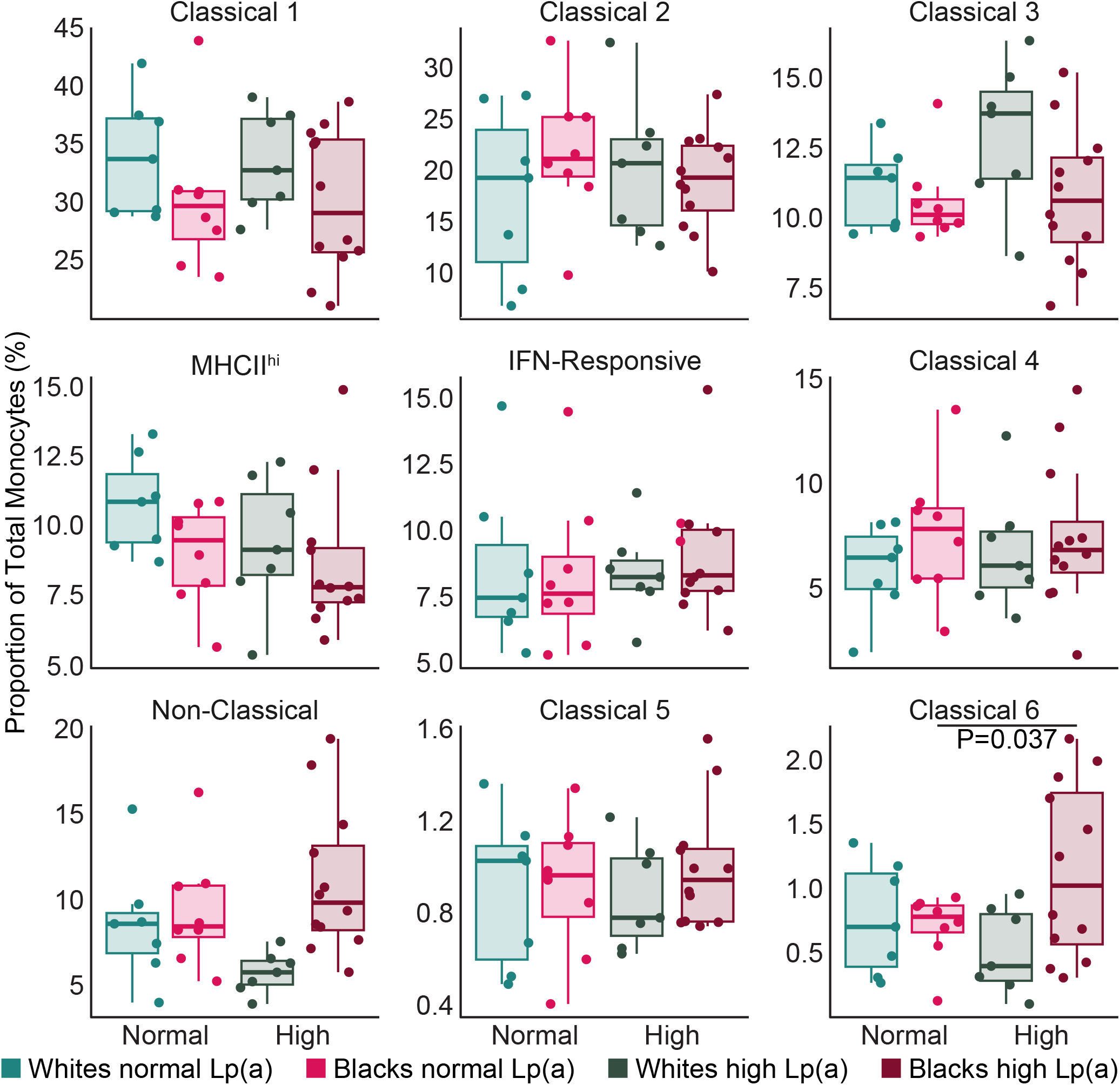
Blacks with high plasma Lp(a) have a higher proportion of Classical 6 monocytes compared to Blacks with normal plasma Lp(a). Boxplots show relationship between plasma Lp(a) category and monocyte subset proportions. Significant *P*-values of the linear regression following centered log-ratio transformation are shown. Results are stratified for race (Normal n_Blacks_=8 and n_Whites_=7; High n_Blacks_=12 and n_Whites_=7). All analyses are adjusted for weighted isoform size and sex.

To further examine these relationships, we performed regression analyses using Lp(a) concentrations as a continuous variable. Lp(a) concentration were log_2_-transformed to estimate changes in monocyte subset proportion for every doubling of plasma Lp(a). These analyses revealed that race modified the association between plasma Lp(a) and non-classical monocytes (*P*_interaction_ = 0.032) (Supplemental Table III). In White participants, plasma Lp(a) concentrations were inversely associated with the proportion of non-classical monocytes, with a 9.2% decrease in this subset for every doubling of plasma Lp(a) after adjusting for wIS and sex (*P* = 0.028) (Figure 3; Supplemental Table III). Similar results were obtained in univariate analysis and after adjustment for wIS only (Supplemental Table III). No such association was observed in Black participants (*P* = 0.470).

**Figure 3.**
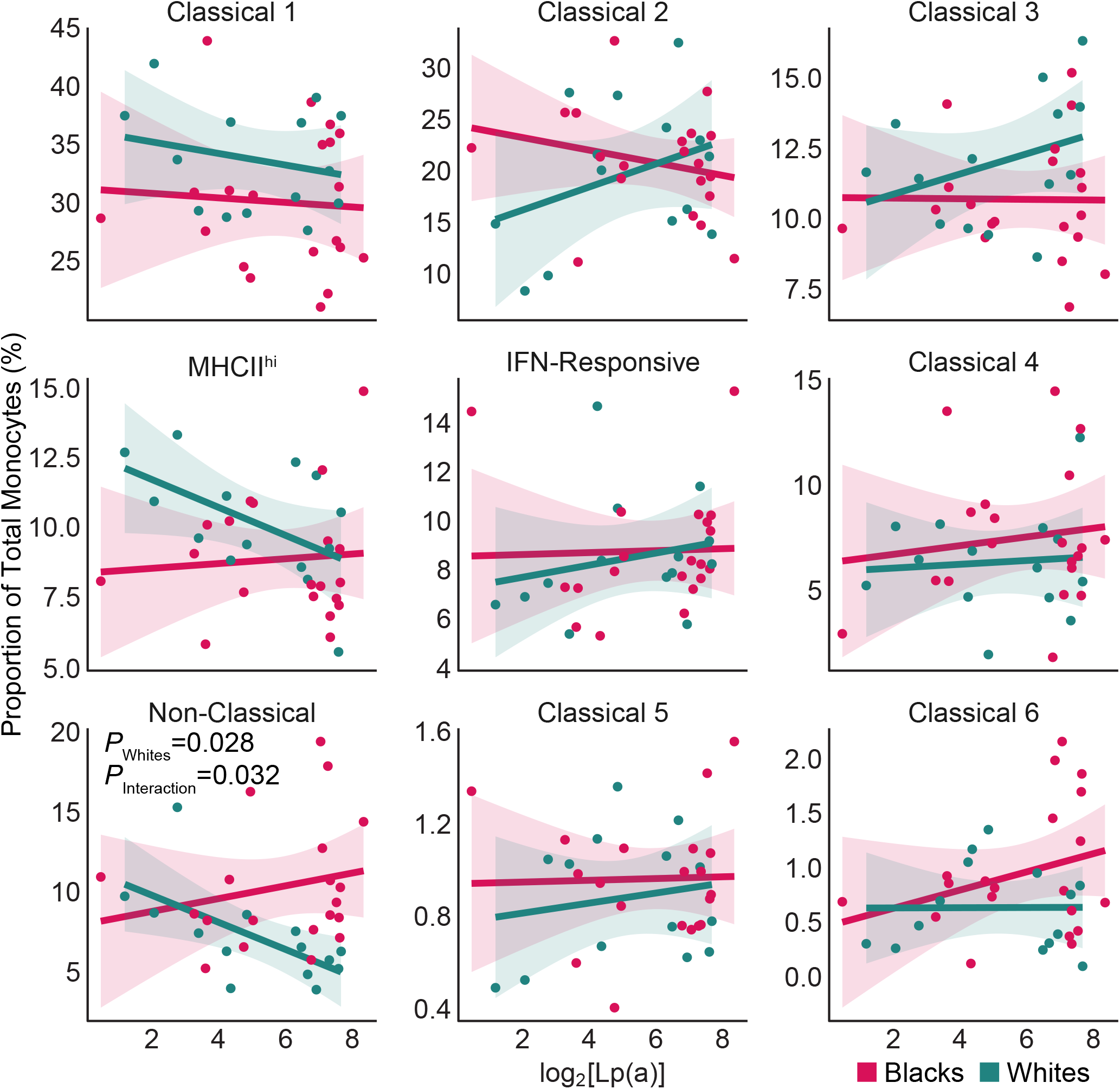
Race moderates the association between plasma Lp(a) concentration and Non-Classical monocytes. Scatterplots show the relationship between log_2_ - transformed plasma Lp(a) levels and monocyte subset proportions. Significant *P*-values of the linear regres-sion using race as an interaction term following centered log-ratio transformation are shown. Results are stratified for Blacks and Whites (n_Blacks_=20; n_Whites_=14). All analyses are adjusted for weighted isoform size and sex.

Race also modified the association of plasma Lp(a) levels with classical 2 monocytes (*P*_interaction_ *=* 0.027). Plasma Lp(a) levels tended to be associated with a higher proportion of classical 2 monocytes in White but not in Black participants (*P*_Whites_ = 0.082; *P*_Blacks_ = 0.172). In addition, plasma Lp(a) showed a trend toward association with higher classical 6 monocytes in Black but not White participants (*P*_Blacks_ = 0.065; *P*_Whites_ = 0.872), although this association was not modified by race (*P*_interaction_ = 0.228). Finally, plasma Lp(a) tended to be negatively associated with MHCII^hi^ monocytes in White participants in univariate models and models adjustment for wIS, but this relationship was no longer significant after adjustment for wIS and sex (Supplemental Table III).

### Oxidized phospholipids bound to APO(a) associate with Lp(a) levels and monocyte subsets

Because oxPL carried by Lp(a) are recognized mediators of its pro-inflammatory effects, we examined whether oxPL were associated with monocyte subset composition in our cohort. Plasma Lp(a) levels were strongly correlated with oxPL bound to APO(a) (R^2^ = 0.84; *P* = 3.3×10^−14^) (Supplemental Figure IIA). Consistent with the association observed with Lp(a), oxPL bound to APO(a) was inversely associated with the proportion of non-classical monocytes in White but not Black participants (*P*_Whites_ = 0.027; *P*_Blacks_ = 0.275; *P*_Interaction_ = 0.014), and tended to be positively associated with classical 2 monocytes in White but not Black participants after adjusting for wIS and sex (*P*_Whites_ = 0.062; *P*_Blacks_ = 0.673; *P*_Interaction_ = 0.079) (Supplemental Table IV).

To determine whether these associations reflected the effects of oxPL more broadly rather than oxPL specifically associated with Lp(a), we examined oxPL bound to APOB. Plasma Lp(a) levels showed a moderate correlation with APOB-bound oxPL (R_2_ = 0.46; *P* = 1.1×10_-5_) (Supplemental Figure IIB). However, APOB-bound oxPL were not associated with the proportions of any monocyte subsets (Supplemental Table V).

Together, these findings indicate that oxPL bound to APO(a) strongly associates with plasma Lp(a) concentrations and show similar race-dependent associations with monocyte subsets, particularly non-classical monocytes. These observations are consistent with the strong correlation between Lp(a) and oxPL carried on APO(a).

## Discussion

In the current study, we leveraged single-cell transcriptomics to characterize the relationship between plasma Lp(a) concentrations and circulating monocyte subsets in White and Black individuals enrolled in a previously completed study. Our analysis suggests that race modifies the association between plasma Lp(a) levels and the proportion of non-classical monocytes. Specifically, plasma Lp(a) levels inversely associate with the proportion of non-classical monocytes in White participants, whereas no association was observed in Black participants. Race also modified the relationship between plasma Lp(a) levels and the proportion of classical 2 monocytes. In addition, Black participants with high plasma Lp(a) levels exhibited a higher proportion of classical 6 monocytes compared to those with normal plasma Lp(a) levels, although this subset represents a relatively small fraction of circulating monocytes and may therefore contribute minimally to disease development. Collectively, these findings suggest race-dependent differences in the relationship between plasma Lp(a) levels and circulating monocyte subpopulation composition.

Previous work from our group and others identified multiple monocyte subsets in healthy individuals and across diverse disease states, including several classical monocyte subsets, non-classical monocytes, MHCII_hi_ monocytes, and IFN-responsive monocytes._19,25–29_ However, the relationship between these subsets and CVD remains incompletely understood. In our previous work, we observed a significant negative association between cigarette smoking and the propotion of non-classical monocytes._19_ The observation of a similar pattern in another CVD risk state, such as high Lp(a), is therefore notable and suggests that alterations in non-classical monocyte abundance may represent a consistent feature of CVD-associated inflammatory states. Interestingly, the inverse relationship between plasma Lp(a) levels and non-classical monocytes was observed in White but not in Black participants. Non-classical monocytes are thought to play a protective role in vascular homeostasis by patrolling the endothelium, exhibiting anti-inflammatory functions, and clearing cellular debris through phagocytosis._30_ Consequently, reduced abundance of this subset has been proposed to increased suspectibility to CVD._31_ However, the role of non-classical monocytes may be context dependent. For example, during viral infections, they can adopt pro-inflammatory functions._31_

Previous work suggests that the association between Lp(a) and CVD is largely driven by oxPL carried on Lp(a). OxPL bound to Lp(a) has been shown to enhance monocyte transendothelial migration *ex vivo* and promote monocyte entry into the arterial wall in humans._17,32_ More recently, diacylglycerol and lysophosphatidic acid bound to Lp(a) have also been implicated in promoting monocyte transendothelial migration *ex vivo*._33_ In our dataset, plasma Lp(a) levels were strongly correlated with APO(a)-bound oxPL, whereas the correlation with APOB-bound oxPL was weaker. Consistent with our observation for plasma Lp(a) alone, race modified the relationship between APO(a)-bound oxPL and non-classical monocytes in White but not Black participants. We attribute this finding to the strong correlation between plasma Lp(a) levels and APO(a)-bound oxPL.

Previous studies identified a positive association between high plasma Lp(a) levels and IFNα and IFNγ responsive gene expression in circulating CD14_+_ monocytes in both healthy individuals and CVD patients._18,34_ Moreover, antisense oligonucleotides targeting APO(a) decreased Lp(a) plasma levels by ~47% and decreased blood IFNα and IFNγ responsive CD14_+_ monocytes._18_ We did not find a relationship between plasma Lp(a) levels and IFN-responsive monocytes. This may be due to the larger relative difference in plasma Lp(a) levels observed in the previous study_18_ compared to our study, the techniques used to identify the monocyte subset, or other population-level differences. Of note, our use of single cell RNA-Seq to analyze the whole blood monocyte population allowed us to detect non-classical monocytes with low CD14 expression that may have been lost in the previous study, which utilized a more direct bulk RNA-seq approach to interrogate CD14_+_ monocytes specifically.

In conclusion, these findings highlight the importance of considering race when examining associations between Lp(a) and circulating monocyte subsets. They also emphasize the need for further studies to elucidate the mechanisms undelrying these differences, including potential *LPA* gene variants, environmental, or epigenetic factors that may influence the interplay between Lp(a) and the innate immune system. Improved understanding of these dynamics may help clarify how Lp(a) contributes to CVD across diverse populations.

### Limitations

This study has several limitations. The analyses were performed using data and stored frozen samples collected as part of a previously published study examining the role of monocytes in cardiovascular disease. Of the 128 individuals enrolled in that study, we restricted our analyses to individuals without diabetes and who were non-smokers in order to minimize potential confounding effects of these conditions on circulating monocyte populations. As a result of these exclusion criteria, the final sample size was limited to 34 individuals. In addition, our analyses were restricted to participants who self-reported race as Black or White, which limits generalizability of these findings to other populations. The modest sample size may also reduce statistical power to detect more subtle associations between Lp(a) levels and additional monocyte subsets. Nevertheless, the use of high-dimensional single-cell transcriptomic profiling provides a unique oppurtunity to examine relationships between Lp(a) and circulating immune cell composition at a level of resolution that is rarely avaiable in larger population-based cohorts.

## Acknowledgements

We would like to thank the research volunteers. We acknowledge the contribution of the Center for Advanced Laboratory Medicine (CALM) at Columbia University Medical Center, the CUIMC Single Cell Analysis Core, Dr. Santica Marcovina (Medpace Inc.) for the Lp(a) and isoform size measurements and, the outpatient unit and Biomarkers laboratory of the Columbia University Irving Institute for Clinical and Translational Research. Oxidized phospholipid measurements were performed by the labs of Drs. Calvin Yeang and Sotirios Tsimikas at The University of Calidfornia San Diego.

